# A DNA G-quadruplex/i-motif hybrid

**DOI:** 10.1101/737296

**Authors:** Betty Chu, Daoning Zhang, Paul J. Paukstelis

**Author notes:** Correspondence: Paul J. Paukstelis.

## Abstract

DNA can form many structures beyond the canonical Watson-Crick double helix. It is now clear that noncanonical structures are present in genomic DNA and have biological functions. G-rich G-quadruplexes and C-rich i-motifs are the most well-characterized noncanonical DNA motifs that have been detected *in vivo* with either proscribed or postulated biological roles. Because of their independent sequence requirements, these structures have largely been considered distinct types of quadruplexes. Here, we describe the crystal structure of the DNA oligonucleotide, d(CCAGGCTGCAA), that self-associates to form a quadruplex structure containing two central antiparallel G-tetrads and six i-motif C-C^+^ base pairs. Solution studies suggest a robust structural motif capable of assembling as a tetramer of individual strands or as a dimer when composed of tandem repeats. This hybrid structure highlights the growing structural diversity of DNA and suggests that biological systems may harbor many functionally important non-duplex structures.

## Introduction

Non-Watson-Crick base pairing interactions in DNA can give rise to a variety of structural motifs beyond the canonical double helix. New types of DNA structural motifs continue to be reported,^1–9^ suggesting that our understanding of DNA’s structural diversity has not been reached. The G-quadruplex and the i-motif are two noncanonical structures that have been studied extensively, and each is characterized by specific types of noncanonical interactions. G-quadruplexes (G4s) are formed from G-rich sequences and contain stacked guanosine tetrads, organized in a cyclic hydrogen bonding arrangement between the Hoogsteen and Watson-Crick faces of neighboring nucleobases.^1, 10^ G4s can be formed through inter- or intramolecular interactions in a variety of topologies and are stabilized by central cations.^11–13^ The DNA i-motif is characterized by the formation of hemiprotonated C-C^+^ parallel-stranded base pairs, which are organized to allow two duplexes to intercalate in an antiparallel fashion to form a quadruplex structure.^2, 14^ Both G4s and i-motifs can form as unimolecular, bimolecular, or tetramolecular assemblies, leading to diverse folding topologies.^15, 16^

Though G4 and i-motif structures tend to form from sequences that contain contiguous stretches of G’s or C’s, respectively, structural characterization has revealed a relatively wide distribution of sequences capable of forming these and similar noncanonical motifs. A unimolecular G4 consensus motif, G_3-5_N_1-7_G_3-5_N_1-7_G_3-5_N_1-7_G_3-5_, was initially used for G4 identification,^17^ leading to initial estimates of ~300,000 possible G-quadruplex forming structures in the human genome.^18^ However, mounting structural evidence indicated that the sequences capable of forming G4s, and the G4 structures themselves, were more diverse than originally thought. Structural variations of G4 structures include motifs that incorporate non-G-tetrads,^19^ bulged residues,^20^ G-triads,^21, 22^ G-tetrads as part of pentad assemblies,^23^ and hybrid G-quadruplex/duplexes.^24, 25^ This sequence and structural diversity led to the doubling of the predicted G4-forming sequences in the human genome to greater than 700,000.^26^ Similarly, a unimolecular i-motif folding rule was formulated based on experimental evidence.^27^ This specified five cytosine residues for each of the four C-tracts, but allowed for greater variation in the length and sequence of the loop regions. Based on this, a preliminary search predicted more than 5000 i-motif-forming sequences in the human genome.^27^ However, isolated i-motif structures with shorter or longer C-tracts have been reported,^28–30^ and the characteristic C-C^+^ base pair of i-motifs is prevalent in a variety of other noncanonical DNA structures,^4, 6, 8, 31, 32^ suggesting that they can serve as building blocks or structural units for other types of structures. Additionally, the structural topology of i-motifs is not limited to only C-C^+^ base pairs. Even the earliest i-motif structures incorporated other noncanonical base pairs^2, 33–36^ or base triples^37, 38^ that stabilize the motif through stacking on the hemiprotonated cytosine base pairs.^39^ As a result, the number of sequences in the human genome with the potential to form i-motifs or related structures is likely much greater than previously predicted.

Both of these noncanonical structural motifs are present in cellular DNA, though their roles in biological processes are just beginning to be understood. G4s have been implicated in a wide variety of normal cellular processes, including DNA replication and transcription, as well as a number of disease states.^40^ Telomeric G4 structures have been visualized using specific antibodies.^41^ The active formation of G4s,^42, 43^ as well as their stabilization by small molecule ligands,^42^ in human cells have also been confirmed. With a predicted 50% of human genes containing G4s at or around promoter regions, DNA G4 structures are predicted to have widespread roles in gene expression.^44^ In particular, the significant enrichment of the G4 motif in a wide range of oncogene promoters suggests its functional importance in cancer.^45^ Examples of G4s modulating gene transcription have been found in the c-MYC,^46^ bcl-2,^47^ and KRAS^48^ oncogene promoters. Additionally, the stabilization of G4s by small molecule ligands at the hTERT^49^ and PDGFR-β^50^ oncogene promoters has been associated with downregulated activity. Nonetheless, the highly thermostable G4s can be detrimental to biological processes and lead to genome instabilities.^40^ DNA i-motifs have long been implicated in biological processes,^27, 45, 51^ but have now been observed *in vivo*. In-cell NMR identified characteristic i-motif signals in HeLa extracts with transfected i-motif DNAs, providing direct evidence that i-motif structures are stable in cellular environments.^52^ Furthermore, the antibody-mediated observation of i-motifs in the nuclei of human cells^53^ and the discovery of i-motif binding proteins that regulate gene activity^54^ demonstrate that i-motifs can have biological function. The sequence and structural diversity of G4s and i-motifs and their growing importance in cellular DNA transactions open the possibility of new variations of these motifs with distinct biological functions.

Here, we describe the crystal structure of a G-quadruplex/i-motif hybrid structure, formed by the oligonucleotide, d(CCAGGCTGCAA). Two distinct strands form a dimer through parallel-stranded interactions, while a symmetry-related dimer interacts in an antiparallel orientation to form the tetramer. The tetramer contains two central G-tetrads that are stabilized by a barium ion, and are flanked on either side by a base triple, one unpaired guanosine, and an i-motif of three C-C^+^ base pairs. Solution studies indicate that the same hybrid quadruplex is formed from tandem sequence repeats, suggesting the potential for this type of structure to form from repetitive DNA elements. d(CCAGGCTGCAA) represents the first structural observation of a G-quadruplex/i-motif hybrid and further expands the wide-ranging structural diversity of DNA.

## Results and Discussion

### Overview

The crystal structure of d(CCAGGCTGCAA) was determined by signal wavelength anomalous dispersion using a 5-Br-deoxyuridine substitution at the T7 position. Initial phases from this derivative were used to create electron density maps for the higher resolution native structure (Table S1). Refined native and derivative structures were virtually identical, with an RMSD of 0.377 Å for all DNA atoms of the asymmetric unit. The asymmetric unit contains two oligonucleotides (Chains A and B) that interact as a dimer. The two monomers show a large degree of structural similarity in the first 5 residues (RMSD, 0.757 Å for 84 atoms), with the largest deviation arising from the differing conformations of the A3 nucleobase (Figure S1a). However, the latter half of the chain contains significant conformational differences in both the backbone and nucleobase atoms (Figure S1b). Two dimers interact through crystal symmetry (symmetry molecules designated as Chains A’ and B’) to form a tetramer. This tetramer contains a number of distinct structural motifs, including a central G-quadruplex, a base triple interaction, a structurally variable spacer region, and a terminal i-motif (Figure 1).

**Figure 1.**
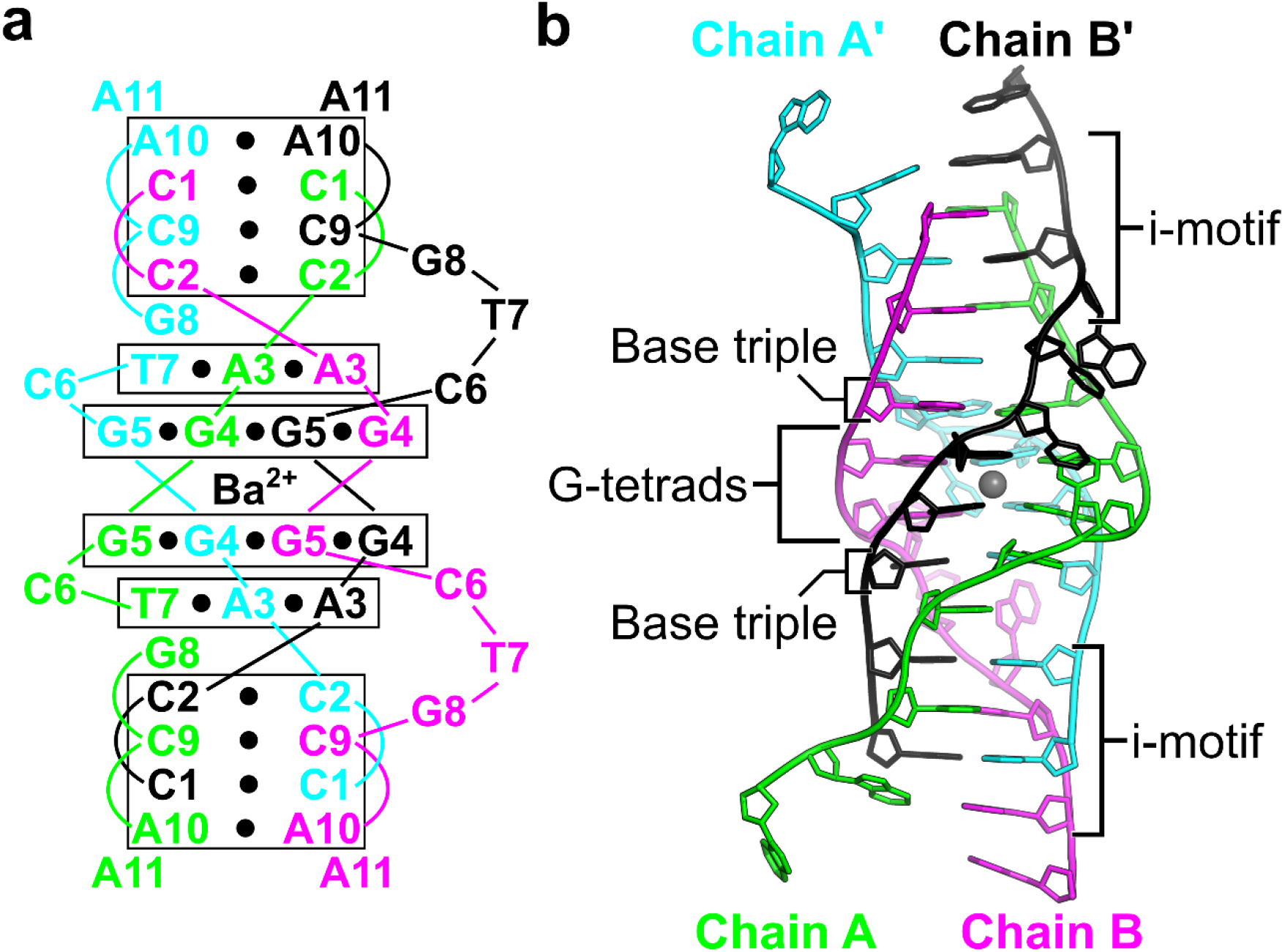
A G-quadruplex/i-motif hybrid structure formed from d(CCAGGCTGCAA). (a) Secondary structure of interactions formed between two symmetry-related dimers. Black circles represent hydrogen bonding interactions. Chains A and A’ are in green and cyan, respectively. Chains B and B’ are in magenta and black, respectively. (b) Cartoon representation of the hybrid quadruplex with labeled features. The gray sphere represents a barium ion.

### A Barium-stabilized G-quadruplex

The central G-quadruplex is composed of two symmetrically equivalent G-tetrads, each of which is formed through two G4 and two G5 residues (Figure 2). The two dimers are antiparallel with respect to each other, with G4-G5 dinucleotide steps along each strand, leading to heteropolar stacking between the two G-tetrads. The G-tetrads are arranged in the *abab* topology.^15, 55, 56^ Like other antiparallel G4s, the tetrad adopts *syn-anti-syn-anti* glycosidic angles with residue G4 in *syn* and G5 in *anti* for both chains. The observed base pair and base step geometries are comparable to other quadruplex structures containing only two G-tetrads with the same topology.^57–59^ This arrangement gives rise to two grooves of distinct widths.^55^ The G5-G5 phosphate distances across the narrow and wide grooves are 12.78 Å and 19.15 Å, respectively.

**Figure 2.**
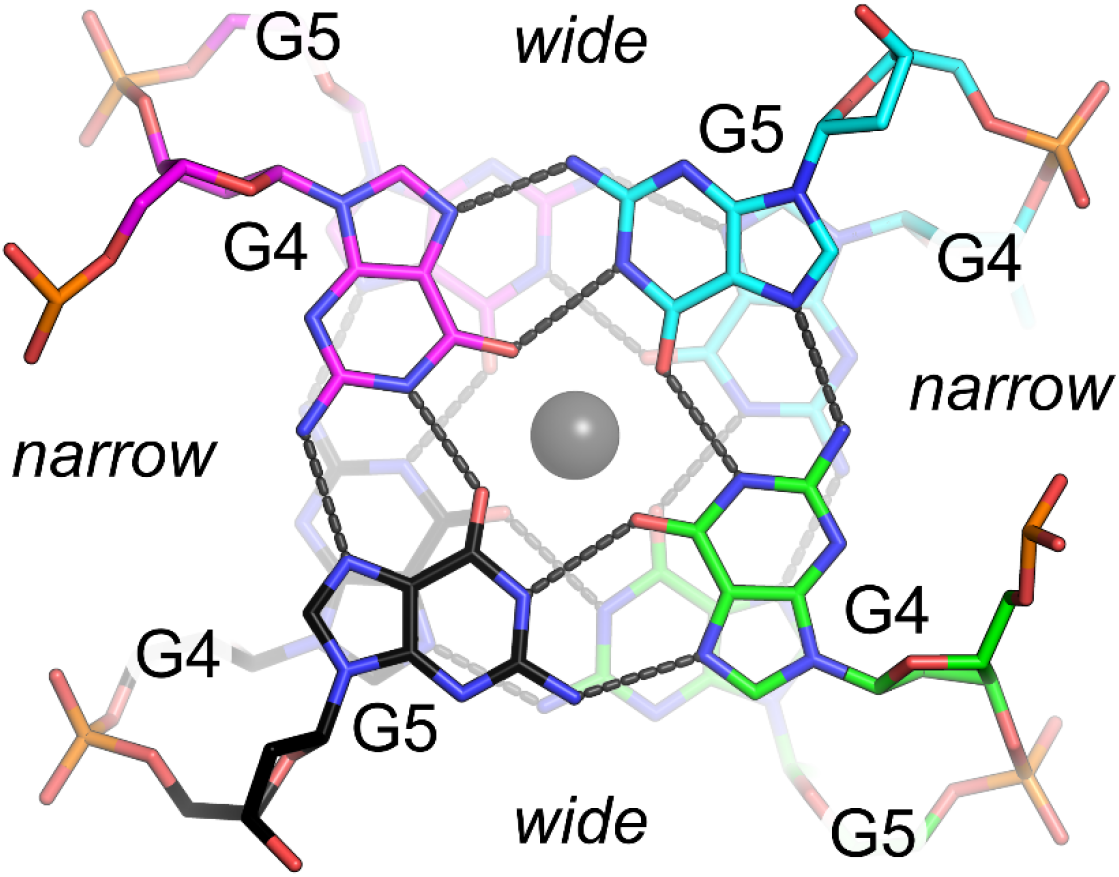
G-quadruplex. Top view of two stacked G-tetrads. Hydrogen bonds are indicated by gray dashes. The gray sphere represents a barium ion. The wide and narrow grooves are indicated.

The eight guanosines coordinate directly with a central cation that is located between the two G-tetrad planes. Both the native and U7-Br oligonucleotides were crystallized in the presence of barium chloride, and a strong (11 σ) anomalous difference electron density peak between the two G-tetrads was observed in both native and derivative structures. This peak is most consistent with Ba^2+^, given the crystallization conditions and data collection energy (Figure S2). The Ba^2+^ ion lies on or very near a crystallographic symmetry axis with a refined final occupancy of 0.50 and B-factors of 73.27 Å^2^. The coordination distances between the cation and guanosine O6 positions range from 2.5 Å to 2.8 Å, with an average distance of 2.63 Å. This average is slightly shorter than the ~2.75 Å average coordination distance observed in previous examples of G-tetrads stabilized by Ba^2+^.^60, 61^ The apparent shorter metal-oxygen coordination may be the result of several factors, including difficulty in refining the cation residing near a special position. Alternatively, this more compact arrangement of guanosine residues could be a structural preference arising from the fewer base stacking interactions on either side of the two G-tetrads.

### Reverse-Hoogsteen Base Triple

Flanking each side of the G-quadruplex is an A-A-T base triple. This noncanonical base triple involves both A3 residues from the dimer and T7 from Chain A’ (Figure 3a). The A3-A3 base pair is formed through the Watson-Crick face of Chain A and the Hoogsteen face of Chain B, which adopts a *syn* glycosidic torsion angle to facilitate the N1-N6 and N6-N7 hydrogen bonds. The base triple is completed by interactions between A3 of Chain A and T7 from Chain A’. This is a reverse Hoogsteen base pair through the N6-O2 and N7-N3 hydrogen bonds. The *syn* glycosidic angle of A3 from Chain B allows the Watson-Crick face to make direct hydrogen bonding contacts with phosphate oxygens of T7 from Chain B’ of the tetramer (Figure 3b). With both N1 and N6 of A3 in hydrogen bonding distance with the non-bridging phosphate oxygens, this arrangement suggests protonation of the N1 position to serve as a hydrogen bond donor. Similar to observations in RNA structures, the electrostatic stabilization between the localized positive charge following N1 protonation and the negatively charged phosphate would facilitate this pKa perturbation.^62^

**Figure 3.**
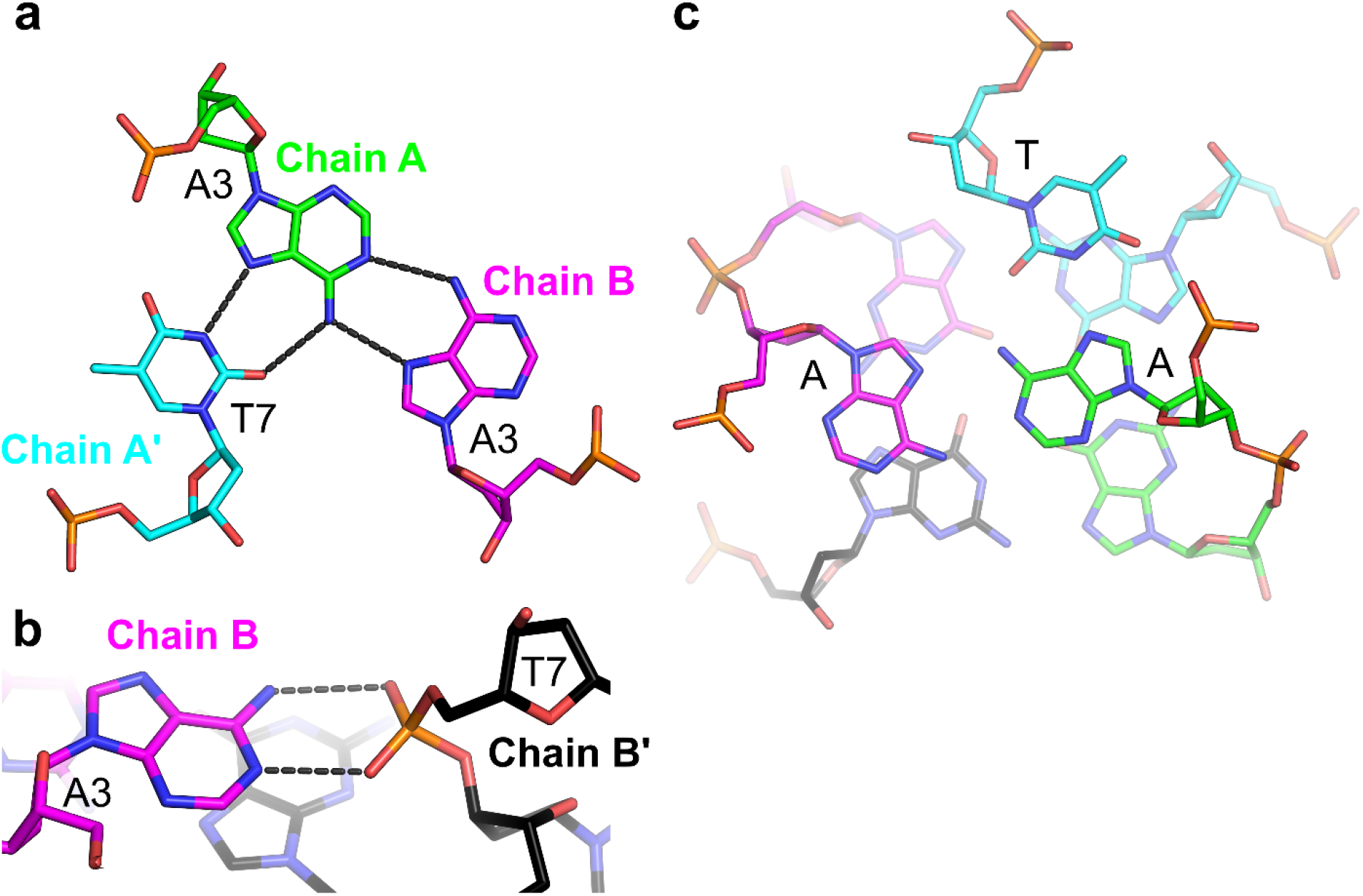
Base triple. (a) Residue A3 from Chain A (green) forms a reverse Hoogsteen base pair with T7 from Chain A’ (cyan) through the A(N6)-T(O2) and A(N7)-T(N3) hydrogen bonds, and interacts with A3 from Chain B (magenta) through the Watson-Crick/Hoogsteen faces. (b) Residue A3 from Chain B hydrogen bonds with the phosphate oxygens of T7 from Chain B’. Hydrogen bonds are indicated by gray dashes. (c) Top view of the A-A-T base triple above one G-tetrad.

Surprisingly, there is little direct nucleobase stacking between the G-tetrad and the A-A-T base triple. Rather, the adenosine and thymidine residues are largely positioned between the tetrad guanosines (Figure 3c, S3a). This is in contrast to the only other example of a base triple flanking one side of a G-quadruplex containing two G-tetrads of the same topology.^63^ In this case, the 22-nucleotide-long d[AGGG(CTAGGG)_3_] contains a C-G-A base triple that forms significant stacking interactions with the G-tetrad (Figure S3b). The large differences in stacking interactions between the triples and the tetrads suggest significant structural variability in these types of interactions based on intrinsic sequence differences and local structural constraints.

### Variable Spacer Region

The most distinct structural differences between the two molecules of the asymmetric unit are in residues C6, T7, and G8 that collectively make up the spacer regions between the central G-tetrad and the peripheral i-motif. Interestingly, these residues have contrasting functional roles in the overall architecture of the tetramer. C6 of Chain A forms a single hydrogen bond with the G5 (N3-N2) from Chain B’ and is tucked into the quartet’s wide groove. C6 in Chain B does not form any base pairing interactions within the tetrameric structure. It is bulged from the tetramer core and serves primarily in mediating crystal contacts through base stacking interactions with the sugar of C6 from Chain A’ and with the nucleobase of A11 from a symmetry-related dimer. As described above, T7 of Chain A is involved in base triple interactions. In contrast, T7 of Chain B is not involved in any base pairing interactions within the tetramer. Instead, this bulged residue base pairs with A11 from a symmetry-related molecule through standard Watson-Crick pairing interactions to stabilize crystal packing. The G8 residues of both molecules are unpaired. In Chain A, G8 is positioned within the nucleobase core, stacking with A3 of the base triple on one face and with the C2-C2^+^ base pair on the other (Figure S4a). However, the G8 residue in Chain B is flipped out from the core, where it stacks with A11 of Chain A from an adjacent symmetry-related molecule to serve in crystal lattice packing contacts (Figure S4b). This stacking is facilitated by A11 adopting a *syn* glycosidic angle, leading to partial stacking of both the pyrimidine and indole rings of the two purines.

These three residues from the parallel-stranded dimer have distinct functions within the structure. In Chain A, they form an integral part of the tetrameric structure, while the same residues in Chain B serve primarily as a bulged spacer that mediates crystal contacts. Because they have the same sequence, either strand could presumably take the role of the structural or bulged strand in solution. Though we cannot rule out the possibility of dynamic switching of these roles within the tetramer, there are several structural clues that suggest that this strand preference may arise at the time of assembly. The base triple interaction provides asymmetry between the parallel strands. This is seen in both the base pairing interactions with T7, and in the *syn* A3 hydrogen bonding interactions with the phosphate from an antiparallel partner. These interactions bring the phosphate toward the stacked tetramer core and bias that partner strand toward bulging its nucleobases outward as found in the spacer. Additionally, the sequestration of the structural T7 in the base triple interaction would strongly bias the following nucleotide, G8, toward being stacked within the tetramer core.

### i-motif and 3-terminal nucleotides

The d(CCAGGCTGCAA) tetramer is capped at either end by i-motifs (Figure 1). The i-motif is comprised of three C-C^+^ base pairs between C1, C2, and C9 residues of the dimers. The terminal C1-C1^+^ base pair gives the i-motif a 5’-E topology.^16^ Residues C1 and C2 of both chains adopt C3’-endo sugar puckers, allowing the sugar-phosphate to stretch to a helical rise of 6.5 Å. This provides the necessary space to allow the C9-C9^+^ base pair from the symmetry-related dimer to intercalate between them (Figure 4a). The geometries of the three hemiprotonated base pairs are similar, with the largest variation in the buckle and propeller angles (Table S2), consistent with what has been observed in other i-motifs.^33, 64^ Complete base pair and base step parameters are listed in Table S2. Like the G-tetrads, the i-motif creates two grooves of dramatically different widths. The wide grooves are generated by the backbones of the parallel base paired strands and the narrow grooves are formed between one parallel-stranded dimer and the intercalated dimer.

**Figure 4.**
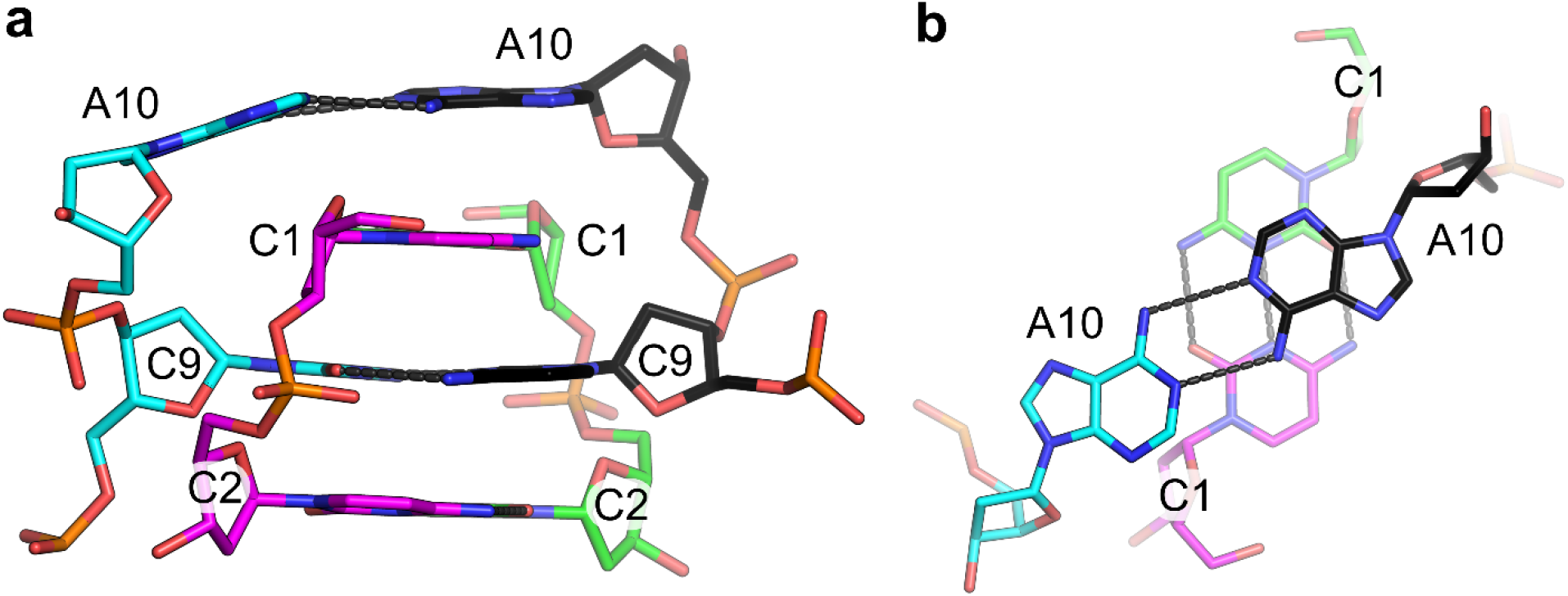
Terminal i-motif. (a) Side view of three C-C^+^ base pairs capped by an A10-A10 base pair. The C9-C9^+^ base pair (cyan/black) is intercalated between the dimeric (green/magenta) C1-C1^+^ and C2-C2^+^ base pairs. (b) Top view of the A10-A10 base pair stacking above the C1-C1^+^ base pair. Hydrogen bonds are indicated by gray dashes.

Along with the C-C^+^ interactions, a noncanonical A10-A10 base pair caps the i-motif. The capping of the 5’-E i-motif by noncanonical (i.e., A-A, T-T) base pairs has been observed in previous examples of i-motif structures.^33, 36, 65^ A large ζ angle between C9 and A10 in Chain A moves residue A10 away from the helical core, preventing direct stacking interactions between A10 and the intercalated C1 (Figure 4b). This creates a strong asymmetry with respect to the neighboring C1-C1^+^ base pair. This asymmetry is likely induced by crystals contacts, most notably those made by the subsequent A11 nucleotides. These A11 residues are not involved in i-motif-like interactions, but form stabilizing contacts with the variable bulged region of another tetrameric assembly (see above).

### NMR Solution Structure Analysis

We conducted 2D-NOESY and 2D-TOCSY experiments on the 11-mer oligonucleotide to directly assess the structure in solution. These spectra suffered from signal crowding and the appearance of multiple conformations that made complete proton assignment difficult. Sequential assignment allowed identification of the nucleobase, H1’, and H2’ protons in at least one conformation. Sugar-to-base connectivities were observed from C2 through G4 (Figure S5) and from C6 through A11 (Figure S6). Four imino proton signals were assigned to T7H3, G4H1, G5H1, and C2H3 (Table S3). These assignments allowed identification of several key structural features.

Three key structural features were confirmed by NMR analysis. First, the C2H3 signal demonstrates protonation at this position and evidence for a C-C^+^ base pair of the i-motif. This was the only CH3 resonance observed. Typically, this proton is observed at chemical shift values near 15 ppm, though in this case there was a significant upfield shift to 12.6 ppm (Figure S7). Cross-peaks to C2H41, C2H42, and C2H5 confirmed the assignment (Figure S7). This large perturbation may be the result of cation-□ interactions, with the localized positive charge at the C2H3 position stacking with the pyrimidine ring of G8 (Figure S4a). NOEs confirmed the proximity of C2H3 and G8, as anticipated from the crystal structure (Figure S7/Table S3). Second, imino NOE cross-peaks confirmed the hydrogen bonding between T7 and A3 and additional NOEs between two independently assigned A3 residues indicated the formation of the A-A-T base triple (Figure S8/Table S3). Third, cross-peaks between the imino protons G4H1 and G5H1 suggest hydrogen bonding between the guanosine residues (Figure S9/Table S3), while resonances between guanosine H8 protons and neighboring guanosine imino and amino protons indicate their interaction through Watson-Crick and Hoogsteen faces. Importantly, these structural features were all internally consistent; NOEs were observed between the guanosines of the tetrads and multiple members of the base triple (Figure S8 (resonances to A3), S9 (resonances to T7)), between the base triple and the unpaired G8 residue (Figure S6), and between G8 and the C2-C2^+^ base pair (Figure S7). Though these NMR data do not allow independent structure determination, they are consistent with the three major base pairing motifs in the crystal structure.

### Tandem repeats alter the oligomeric solution state

The crystal structure suggested that the flexibility at the A11 position could allow tandem sequence repeats to form a dimeric quadruplex, analogous to loops in bi- and unimolecular G4 and i-motif forming sequences. We synthesized the 22-mer tandem repeat, d(CCAGGCTGCAACCAGGCTGCAA), and compared it to the 11-mer by circular dichroism (CD) and UV absorption spectroscopy. The results demonstrate that the assemblies are structurally similar, that the dimeric assembly is significantly more stable than the tetramer, and that both are preferentially stabilized by Ba^2+^.

CD spectra of the 11-mer oligonucleotide titrated with Ba^2+^ (up to 100 mM) showed the appearance of a positive band at ~240 nm, a strong negative band at ~255 nm, and a weak negative band at ~295 nm (Figure 5a). CD melting analysis suggested that these features were due to the formation of a specific structure, with nearly identical forward and reverse temperature dependence spectra (Figure S10a). The 22-mer had a similar CD profile with more pronounced characteristic peaks, suggesting that the tandem repeat forms the same or similar structure as that of the 11-mer (Figure 5b). UV melting analysis showed a dramatic difference in melting temperature between these two assemblies: 41.7±1.3°C for the tetrameric assembly and 73.7±2.5°C for the dimeric assembly (Figure 5c). The fewer number of DNA strands in the dimeric quadruplex likely results in reduced end fraying, which can account for the apparent stability increase in melting experiments and the strength of CD signals.

**Figure 5.**
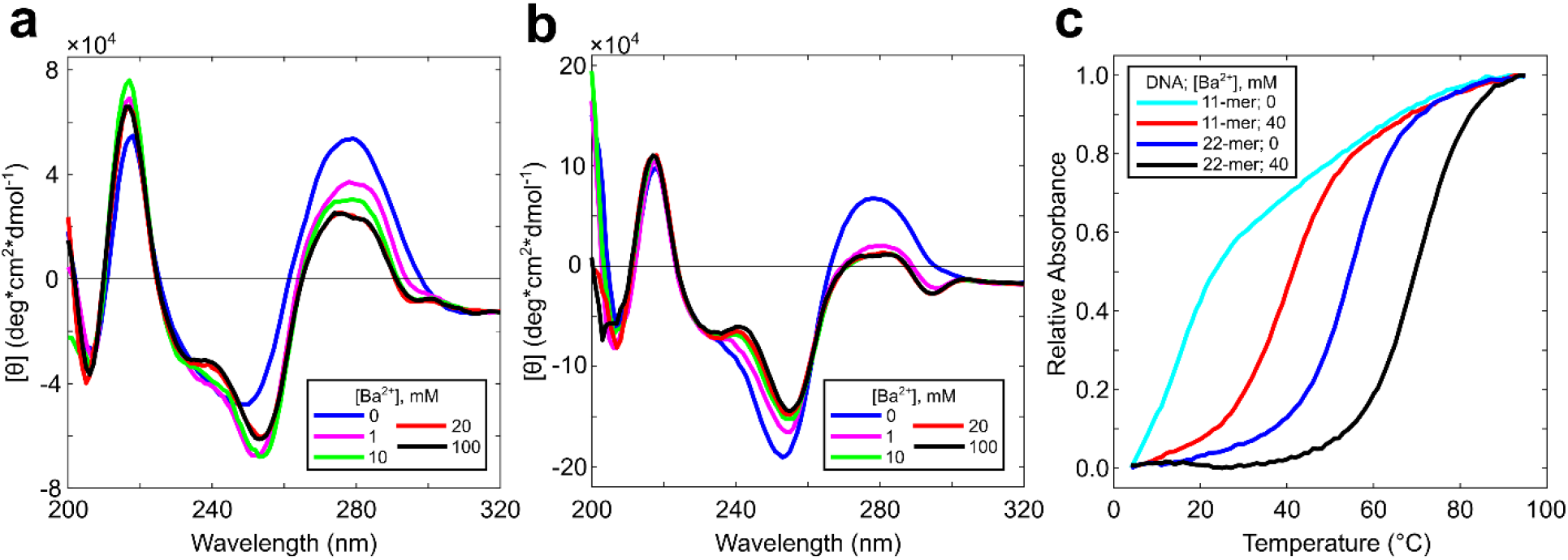
CD and Absorbance spectra. Ba^2+^ titration up to 100 mM at room temperature for (a) the 11-mer, d(CCAGGCTGCAA), and (b) the 22-mer, d(CCAGGCTGCAACCAGGCTGCAA). (c) Thermal denaturation curves from 4°C to 95°C of the 11-mer and 22-mer in sodium cacodylate buffer pH 6.0 or supplemented with 40 mM Ba^2+^.

Both the tetrameric and dimeric assemblies showed a preference for Ba^2+^ over monovalent cations. The spectra for both sequence lengths differed slightly in monovalent cations (K^+^ or Na^+^), with respect to Ba^2+^, showing a shoulder at ~240 nm and a negative band at ~255 nm, but lacking the ~295 nm negative band (Figure S10b). These bands were largely absent for the 11-mer in buffer alone, suggesting additional cations were necessary for structure formation. Thermal denaturation experiments of the 11-mer showed no observable melting transition in conditions containing the buffer alone, whereas melting transitions were observed with additional K^+^ or Na^+^ (37.2±3.9°C and 37.3±4.1°C, respectively; Figure S10c). The 22-mer showed comparable CD profiles between 100 mM monovalent conditions and buffer only, suggesting that the Na^+^ cation from the cacodylate buffer was sufficient to induce some assembly. A melting transition at 56.7±0.3°C was observed in buffer, while additional K^+^ or Na^+^ increased the T_m_ (62.9±0.2°C and 65.3±1.3°C, respectively; Figure S10d). For both the 11-mer and 22-mer oligonucleotides, the observed T_m_ values in monovalent conditions were lower than that in Ba^2+^, suggesting that the divalent cation plays a significant role in structural stability.

### An Alternative Hybrid motif

Finally, we determined that d(CCAGGCTGCAA) can also assemble into an alternative hybrid structure. We crystallized a 5-Br-deoxycytidine substitution at the C9 position and determined its structure (Table S1). The bromine substitution and different crystallization conditions resulted in an overall different structure, but with some similar features. The two molecules in the asymmetric unit interact with symmetry-related strands to form a hybrid quadruplex structure, in this case juxtaposing an i-motif at the 5’ end and a partial antiparallel duplex at the 3’ end (Figure 6).

**Figure 6.**
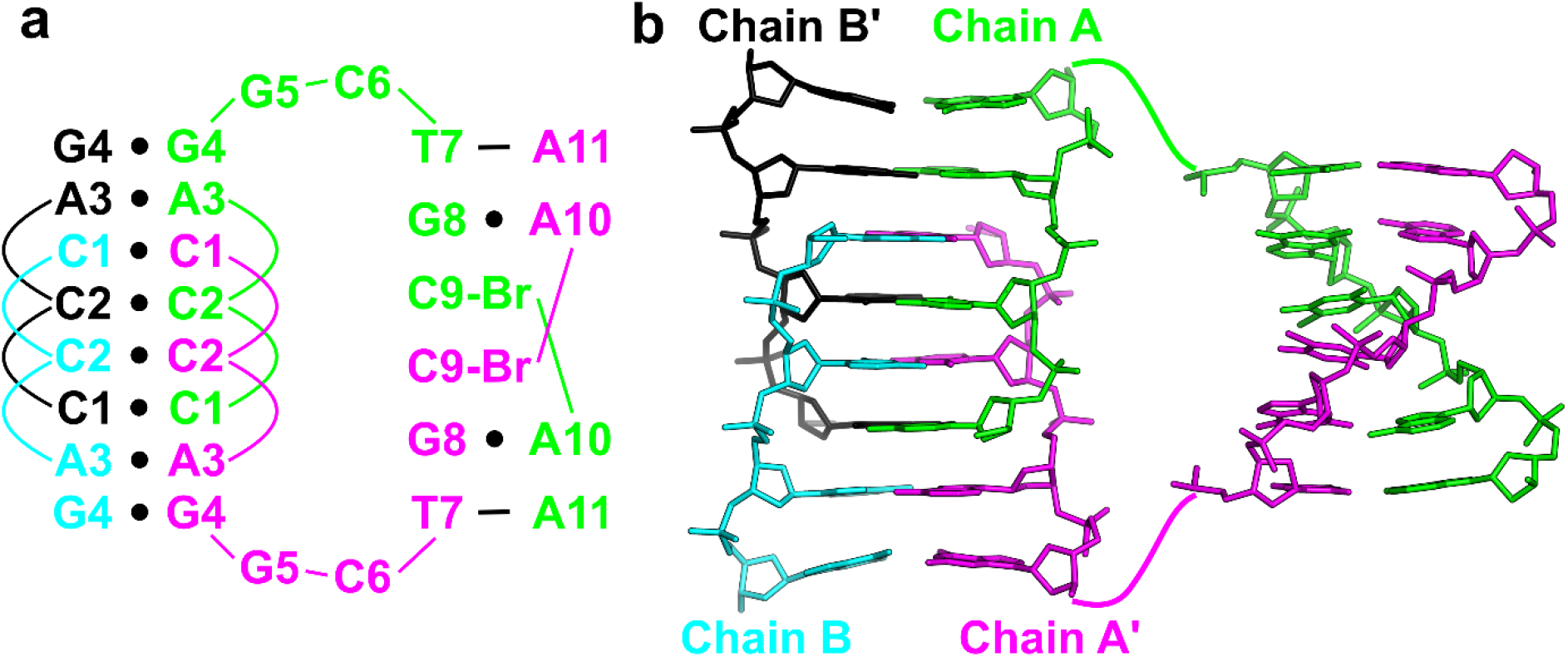
An i-motif/duplex hybrid structure formed from the C9-Br derivative, d(CCAGGCTGC^Br^AA). (a) Secondary structure representation of interactions formed between symmetry-related strands. Black dashes represent Watson-Crick base pairs. Black circles represent noncanonical base pairs. (b) Stick representation of the 5’ i-motif-consisting CCAG homo base stacking motif linked to the 3’ antiparallel duplex. The i-motif and duplex are rotated 70° about the x-axis with respect to each other. Chains A and A’ are in green and magenta, respectively. Chains B and B’ are in cyan and black, respectively.

In this structure, the i-motif C-C^+^ base pairs are formed exclusively from residues C1 and C2. The four strands create a 5’-E topology, with the symmetry axis between the intercalated C2-C2^+^ base pairs. This i-motif region is extended on either side by a homo base pairing region that includes symmetric A3-A3 (N6-N7) and G4-G4 (N1-O6) pairs. The base stacking interactions provided by these noncanonical base pairs stabilize the i-motif tertiary interactions. The brominated C9 residues are no longer involved in i-motif formation, and instead form Watson-Crick base pairing interactions with G5 from symmetry-related molecules that promote crystal packing. Examination of the native structure suggests that the C9 bromine substitution would preclude formation of the hybrid G4/i-motif structure due to significant steric clashes between the bromine in Chain B and the phosphodiester backbone.

The 3’ end of the structure is a short imperfect duplex with noncanonical features. Symmetry interactions between two identical strands (Chains A and A’) form an antiparallel base pairing arrangement consisting of the brominated C9 residues at the interior, each of which is immediately flanked by a G8-A10 base pair formed through N2-N7 and N3-N6 hydrogen bonding, and finally capped by a T7-A11 Watson-Crick base pair (Figure S11). This duplex interacts with a second duplex that is formed from the other unique molecule (Chains B and B’) in the asymmetric unit. The two distinct duplexes are held together by the G5-C9-Br base pair described above and by the C-G-A base triple, which is converted from the G8-A10 base pair from a single duplex (Figure S12).

### Structural and Biological Implications

Though previous biophysical studies characterized oligonucleotides capable of forming a parallel G4/i-motif hybrid in solution,^66^ the results presented here provides the first structural snapshot of a hybrid G4/i-motif. This information provides the beginnings of a structural paradigm for how these two distinct quadruplex motifs can coexist. Most notably, this structure suggests a requirement for spacer elements to separate the two base pairing motifs. These spacer elements serve to bridge the large differences in interstrand backbone distances of the two motifs. This is necessitated by an exchange of the wide and narrow grooves between the individual motifs; the G-tetrad wide groove is continuous with the i-motif narrow groove and vice versa. The variable spacer regions that include the structurally integrated base triple and unpaired guanosine allow progressive changes of interstrand backbone distances to facilitate this transition.

Biologically, this structure hints at the potential complexity of noncanonical DNA structures that may be harbored within genomes. The demonstration that the DNA studied here forms a highly stable dimeric structure from tandem repeats suggests that longer repetitive sequences may have the ability to form complex structures, perhaps containing existing known DNA motifs. Repetitive DNA makes up more than 50% of the human genome,^67^ with microsatellite (1-10 nt), minisatellite (10 to several hundred nt), and macrosatellite (up to thousands of nt) repeats, making up ~3%.^68^ Satellite DNA is involved in a variety of biological functions and pathologies,^69^ and repeat sequences have been implicated as drivers of evolution through the formation of noncanonical structures that result in genomic instability.^70^ Though there are now some examples for how repetitive DNA can impact biological function, the structural basis for this is largely unknown. The discovery of new, stable noncanonical DNA structures suggests the possibility that these repeat sequences can form complex motifs that may not be predictable from existing sequence/structure relationships.

## Experimental Section

### Oligonucleotide Synthesis and Purification

d(CCAGGCTGCAA), d(CCAGGCU^Br^GCAA), and d(CCAGGCTGC^Br^AA) were synthesized using standard phosphoramidite chemistry on an Expedite 8909 Nucleic Acid Synthesizer (PerSeptive Biosystems, Inc.), with reagents from Glen Research (Sterling, VA). DNA oligonucleotides were purified using the Glen-Pak cartridges according to the manufacturer’s protocol.

### Crystallization and Data Collection

Sitting drops of d(CCAGGCTGCAA) were set up by mixing 1 μL of 500 μM DNA solution with 2 μL of crystallization solution (30% polyethylene glycol 400 (PEG400), 20 mM barium chloride, 10 mM spermidine, and 30 mM Bis-Tris at pH 8.5). Sitting drops of d(CCAGGCU^Br^GCAA) were set up by mixing 1 μL of 500 μM DNA solution with 2 μL of crystallization solution (25% polyethylene glycol 400 (PEG400), 40 mM barium chloride, 10 mM spermidine, and 30 mM Bis-Tris at pH 8.5). These drops were equilibrated against 300 μL of 5% PEG400 in the well reservoir at 22°C for 15–20 h, followed by subsequent equilibration with 3–4 μL of glacial acetic acid added to the well reservoir. Crystals were observed 2 days after the addition of acid. Crystals were removed from the drops by nylon cryoloops and directly cryo-cooled in liquid nitrogen.

d(CCAGGCTGC^Br^AA) was crystallized by mixing 3 μL of 500 μM DNA solution with 3 μL of crystallization solution (15% 2-methyl-2,4-pentanediol (MPD), 120 mM calcium chloride, 20 mM lithium chloride, 8 mM spermidine, and 30 mM sodium cacodylate at pH 5.5). Crystallization was performed at 22°C and in sitting drops, which were equilibrated against 300 μL of 20% MPD in the well reservoir. Crystals were observed 2 days after plating. Crystals were removed from the drops by nylon cryoloops, dipped in 30% MPD and cryo-cooled in liquid nitrogen.

Diffraction data for d(CCAGGCTGCAA) and d(CCAGGCU^Br^GCAA) were collected at the Advanced Photon Source (APS) 24-ID-C. Diffraction data for d(CCAGGCTGC^Br^AA) were collected at APS 22-BM.

### Structure Determination

Data processing for d(CCAGGCTGCAA) and the U7-Br derivative, d(CCAGGCU^Br^GCAA), was carried out in XDS^71^ and Aimless.^72, 73^ Diffraction data for the C9-Br derivative, d(CCAGGCTGC^Br^AA), were indexed and integrated using iMosflm.^74^ In both derivative datasets, initial phases were determined by single-wavelength anomalous dispersion (SAD) phasing, using SHELX^75^ in CCP4i2.^76^ Two bromine sites were identified in each map, which enabled model building of two chains of each derivative in Coot.^77^ Subsequent refinement was carried out in Refmac.^78, 79^ The refined U7-Br derivative structure was used as a molecular replacement search model in Phaser^80^ for the native oligonucleotide. Further refinement was carried out in Refmac and additional model building was performed in Coot. The PDB-REDO^81^ web server was used to conduct *k*-fold cross-validation of R_free_ values on all three structures and to generate the final models. Final refinement statistics are shown in Table S1. Coordinates have been deposited under the PDB accession codes 6TZQ, 6TZR, and 6TZS.

### Nuclear Magnetic Resonance (NMR) Spectroscopy

NMR data were acquired on a Bruker Avance III 600-MHz spectrometer equipped with a Cryo-TCI probe. The 11-mer oligonucleotide was prepared at 500 μM in 30 mM sodium cacodylate buffer at pH 6.0 containing 40 mM BaCl_2_. A combination of 2D-NOESY and 2D-TOCSY experiments were performed at 10°C, in which the mixing time was set to 100 ms for the 2D-NOESY and 90 ms for the 2D-TOCSY. The oligonucleotide sequential assignment was conducted using the Computer Aided Resonance Assignment (CARA) program.^82^

### Circular Dichroism (CD) Spectroscopy

CD spectra were acquired using the Jasco J-810 spectropolarimeter fitted with a thermostated cell holder. Samples were prepared in 30 mM sodium cacodylate buffer at pH 6.0 containing varying concentrations of BaCl_2_ or 100 mM monovalent (KCl or NaCl) cations. The 11-mer and 22-mer oligonucleotides were prepared at final DNA concentrations of 100 μM and 75 μM, respectively. Samples were equilibrated for 12–18 h at 4°C prior to the acquisition of the spectra. All spectra were collected at room temperature from 200 to 320 nm with a data pitch of 1.0 nm. For melting experiments, the sample was allowed to dwell for 7 min at the temperature set point.

### Thermal Denaturation

UV melting spectra were acquired using the Cary100 Bio UV-visible spectrophotometer equipped with a 12-cell sample changer and a Peltier heating/cooling system. The sample chamber was purged with N_2_ throughout both melting and annealing data collection runs. Samples were prepared in 30 mM sodium cacodylate buffer at pH 6.0 supplemented with 40 mM BaCl_2_ or 100 mM monovalent (KCl or NaCl) cations. The 11-mer and 22-mer oligonucleotides were prepared at final DNA concentrations of 14.4 μM and 7.25 μM, respectively. Samples were equilibrated for 15–20 h at 4°C prior to the acquisition of the spectra. Samples were transferred to self-masking quartz cuvettes with 1 cm path length for UV absorbance measurements. All spectra were collected at 260 nm. An initial fast heating ramp from 4°C to 95°C at 10°C/min was done. Data were collected every 1°C during a slow cooling ramp from 95°C to 4°C at 1°C/min and a subsequent slow heating ramp at the same temperature range and rate. Thermal melting analyses and curve fitting were conducted using MATLAB.^83^

## Acknowledgements

We thank the staff at Northeastern Collaborative Access Team (NE-CAT) and the Southeast Regional Collaborative Access Team (SER-CAT) at the Advanced Photon Source (APS) for their assistance with X-ray beamlines. We thank Dr. Jason Kahn for the use of the UV-Vis spectrophotometer and Dr. Dorothy Beckett for insightful discussions. This work is based upon research conducted at the NE-CAT beamlines, which are funded by the National Institute of General Medical Sciences from the National Institutes of Health (P30 GM124165) and SER-CAT beamlines, which are supported by grants (S10_RR25528 and S10_RR028976) from the National Institutes of Health. This research used resources of APS, a U.S. Department of Energy (DOE) Office of Science User Facility, operated for the DOE Office of Science by Argonne National Laboratory.

## Notes

The authors declare no competing financial interest.

