## Supplementary Material for "A DNA G-quadruplex/i-motif hybrid"

Correspondence:

**Table S1.** Data Collection and Refinement Statistics.*

| | **Native** | **U7-Br Derivative** | **C9-Br Derivative** |
|---|---|---|---|
| **PDB ID** | 6TZQ | 6TZR | 6TZS |
| **Sequence** | d(CCAGGCTGCAA) | d(CCAGGCU$^{Br}$GCAA) | d(CCAGGCTGC$^{Br}$AA) |
| **Beamline** | NE-CAT 24-ID-C | NE-CAT 24-ID-C | SER-CAT BM22 |
| **Data Collection** | | | |
| Space Group | $P3_221$ | $P3_221$ | $I4_122$ |
| Cell Dimensions | | | |
| a, b, c (Å) | 37.38, 37.38, 98.65 | 37.10, 37.10, 98.31 | 51.52, 51.52, 112.95 |
| α, β, γ (°) | 90, 90, 120 | 90, 90, 120 | 90, 90, 120 |
| Resolution (Å) | 49.33 – 2.29 (2.37 – 2.29) | 32.77 – 2.40 (2.49 – 2.40) | 55.80 – 2.60 |
| $R_{meas}$ (within I+/I-) | 0.142 (1.708) | 0.083 (0.917) | 0.177 (1.889) |
| $R_{meas}$ (all I+ and I-) | 0.142 (1.702) | 0.090 (0.917) | 0.182 (1.895) |
| $R_{pim}$ (within I+/I-) | 0.045 (0.543) | 0.037 (0.399) | 0.072 (0.712) |
| $R_{pim}$ (all I+ and I-) | 0.035 (0.400) | 0.032 (0.298) | 0.0058 (0.526) |
| No. of unique | 3955 (373) | 3399 (358) | 2540 (294) |
| I / σ I | 8.2 (1.4) | 11.0 (1.8) | 11.5 (1.2) |
| Completeness (%) | 99.9 (100.0) | 99.9 (100.0) | 100.0 (100.0) |
| Multiplicity | 17.4 (18.0) | 8.9 (9.4) | 12.2 (12.7) |
| Wavelength (Å) | 0.9196 | 0.9196 | 0.9187 |
| **Phasing** | | | |
| Atom/Sites | | Br/2 | Br/2 |
| CFOM from SHELX[1] | | 72.7 | 84.4 |
| **Refinement** | | | |
| Resolution (Å) | 32.88 – 2.29 (2.35 – 2.29) | 32.15 – 2.40 (2.46 – 2.40) | 46.78 – 2.60 (2.67 – 2.60) |
| No. reflections | 3560 | 3040 | 2272 |
| No. reflections used in $R_{free}$ Test Set | 373 | 338 | 254 |
| $R_{work}$** | 0.2223 | 0.2121 | 0.2665 |
| $R_{free}$** | 0.2551 | 0.2538 | 0.3165 |
| $R_{complete}$** | 0.2551 | 0.2516 | 0.3156 |
| Total No. of atoms | 446 | 445 | 453 |
| Average B-factors (Å$^2$) | 71.434 | 79.866 | 46.873 |
| RMS deviations | | | |
| Bond lengths (Å) | 0.0055 | 0.0062 | 0.0076 |
| Bond angles (°) | 1.5652 | 1.6400 | 1.6148 |

*Values in parentheses correspond to the high-resolution shell.

**$R_{work}$, $R_{free}$, and $R_{complete}$ values are from 10-fold cross-validation in PDB-REDO.[2]

**Table S2.** Base pair and base pair step parameters, calculated with x3DNA-DSSR,[3] for the helical core region of the d(CCAGGCTGCAA) tetramer.

***Local Base Pair Parameters***

| | Base Pair | Shear | Stretch | Stagger | Buckle | Propeller | Opening |
|---|---|---|---|---|---|---|---|
| ***1*** | C1-C1 | 1.97 | 1.50 | -0.04 | -2.24 | -0.02 | 177.33 |
| ***2*** | C9-C9 | 2.09 | 1.33 | -0.01 | -2.36 | 6.52 | 177.39 |
| ***3*** | C2-C2 | 2.04 | 1.44 | 0.16 | -0.24 | -3.42 | 179.23 |
| ***4*** | A3-A3 | -4.16 | 1.39 | 0.69 | 8.05 | 10.39 | -112.03 |
| ***5*** | G4-G5 | -1.45 | -3.52 | 0.55 | -16.11 | 10.74 | 87.68 |
| ***6*** | G5-G4 | 1.65 | 3.38 | -0.05 | 8.71 | 0.68 | -89.11 |
| ***7*** | A3-A3 | -4.16 | 1.39 | 0.69 | 8.05 | 10.40 | -112.04 |
| ***8*** | C2-C2 | 2.04 | 1.44 | 0.16 | -0.24 | -3.42 | 179.23 |
| ***9*** | C9-C9 | 2.09 | 1.33 | -0.01 | -2.36 | 6.51 | 177.40 |
| ***10*** | C1-C1 | 1.97 | 1.50 | -0.04 | -2.25 | -0.02 | 177.32 |

***Local Base Pair Step Parameters***

| | Step | Shift | Slide | Rise | Tilt | Roll | Twist |
|---|---|---|---|---|---|---|---|
| ***1*** | CC/CC | -2.48 | 2.23 | 0.09 | 126.28 | 126.29 | -136.14 |
| ***2*** | CC/CC | 1.70 | -2.59 | 0.02 | 148.83 | 95.33 | -82.42 |
| ***3*** | CA/AC | -3.99 | -5.66 | 1.62 | -169.56 | 54.42 | 21.15 |
| ***4*** | AG/GA | 1.98 | 3.58 | -3.52 | -0.89 | -2.11 | -95.65 |
| ***5*** | GG/GG | -2.39 | -2.26 | -1.95 | 86.27 | -155.46 | 39.34 |
| ***6*** | GA/AG | -0.05 | 3.29 | 3.15 | -1.71 | -0.38 | -85.84 |
| ***7*** | AC/CA | 3.99 | 5.66 | -1.62 | 169.56 | -54.41 | -21.14 |
| ***8*** | CC/CC | -1.70 | 2.59 | -0.02 | -148.82 | -95.33 | 82.37 |
| ***9*** | CC/CC | 2.48 | -2.23 | -0.08 | -126.28 | -126.28 | 136.23 |

**Table S3.** Chemical shift values (in ppm) of $^1$H assignments obtained from 2D-NOESY/TOCSY NMR spectra of the 11-mer, d(CCAGGCTGCAA).

| | *Imino* | *Amino* | *H8/H6* | *H5/Methyl* | *H1'* | **H2', H2''* | *H3'* | *H4'* | **H5', H5''* |
|---|---|---|---|---|---|---|---|---|---|
| ***C1*** | - | | 7.39 | 5.53 | 5.75 | 1.73, 1.66 | 4.34 | 3.89 | 3.48, 3.43 |
| ***C2*** | 12.66 | 8.23, 6.85 | 7.40 | 5.65 | | 2.05, 1.90 | 4.69 | | |
| ***A3a*** | - | 7.56, 7.49 | | | | | | | |
| ***A3b*** | - | 10.21, - | 8.04 | - | 5.63 | 2.54, 2.62 | 4.79 | | |
| ***G4*** | 12.89 | 7.45, 7.39 | 7.61 | - | 5.74 | 2.29, 2.62 | 4.79 | | |
| ***G5*** | 12.72 | 7.62, 7.48 | 7.62 | - | 5.58 | 2.28, 2.61 | 4.66 | | |
| ***C6*** | - | 6.54, 6.25 | 7.22 | 4.97 | 5.87 | 1.80, 2.22 | 4.54 | 4.08 | |
| ***T7*** | 13.83 | - | 7.03 | 1.41 | 5.52 | 1.65, 2.00 | 4.61 | 3.86 | |
| ***G8*** | - | 5.87, 5.66 | 7.63 | - | 5.71 | 2.26, 2.34 | 4.66 | | |
| ***C9*** | - | | 7.19 | 5.58 | 5.66 | 1.49, 1.95 | 4.44 | 3.78 | |
| ***A10*** | - | | 7.86 | - | 5.73 | 2.21, 2.35 | 4.63 | 3.99 | 3.74, 3.66 |
| ***A11*** | - | | 7.98 | - | 5.96 | 2.20, 2.40 | 4.47 | 3.93 | |

*H2'/H2'' and H5'/H5'' protons were not stereospecifically assigned.

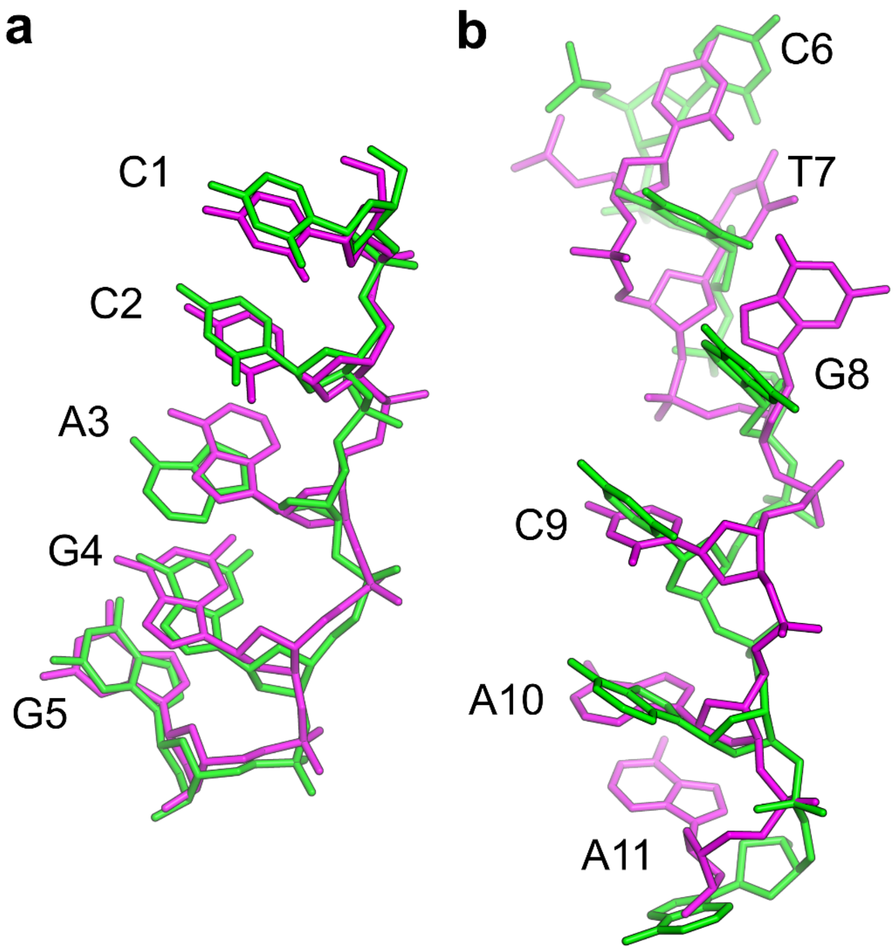

**Figure S1.** Structural Comparison of d(CCAGGCTGCAA) Monomers. Stick representation of the alignment between Chains A (green) and B (magenta) reveal structural similarity in residues 1-5 (a) and significant differences in residues 6-11 (b).

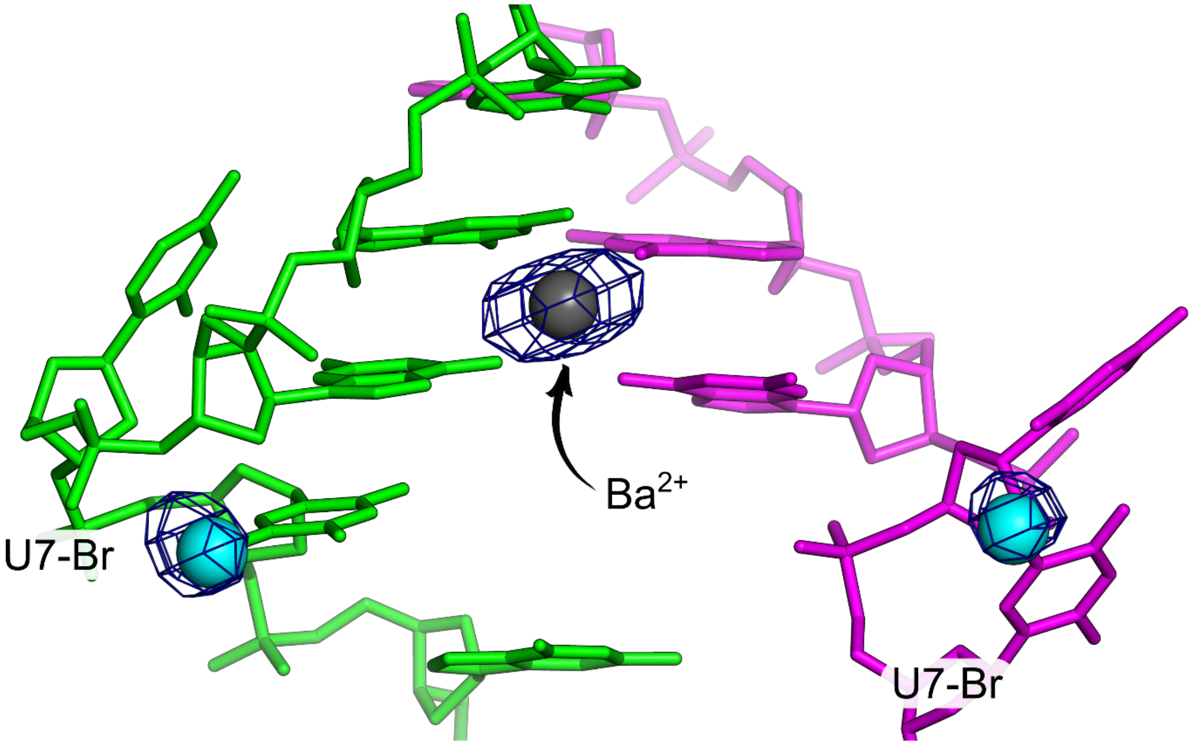

**Figure S2.** Anomalous Differences of the U7-Br Derivative. Anomalous difference electron density (dark blue) contoured at 6.2 σ corresponds to the bromine atoms (cyan spheres) used for phasing. A $Ba^{2+}$ ion is shown as a gray sphere in anomalous electron density.

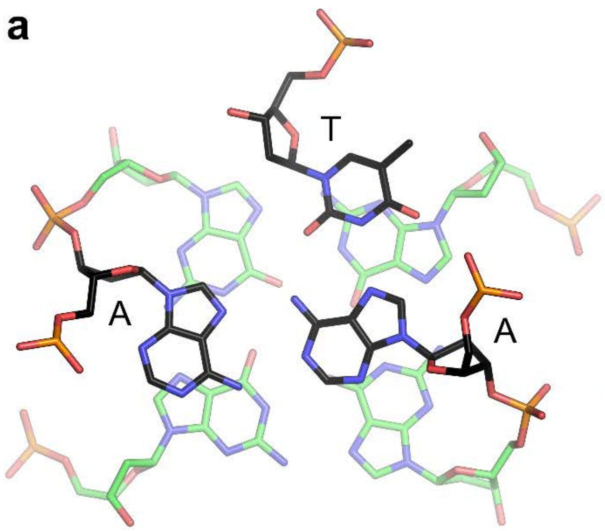

**Figure S3.** Structural Comparison of the Base triple/G-tetrad Stacking between the 11-mer and 2KM3. A base triple (black) stacks above a G-tetrad (green) with the *abab* topology of guanosine residues in (a) the 11-mer and (b) PDB 2KM3.[4]

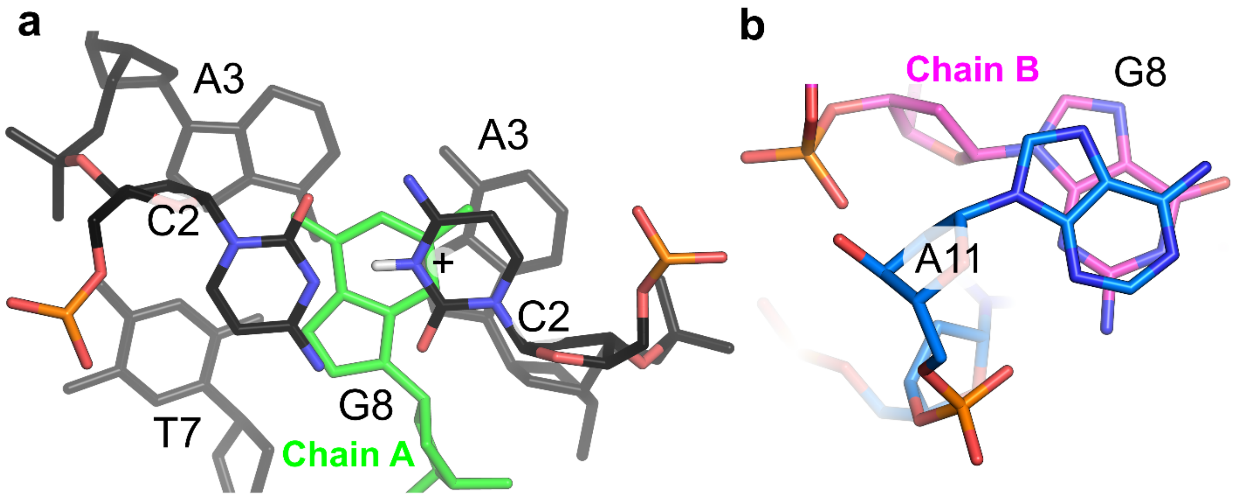

**Figure S4.** G8 Interactions. (a) The unpaired G8 from Chain A (green) stacks between the C2-C2$^{+}$ base pair and the A-A-T base triple. (b) The bulged G8 from Chain B (magenta) stacks with A11 from a symmetry-related molecule (blue).

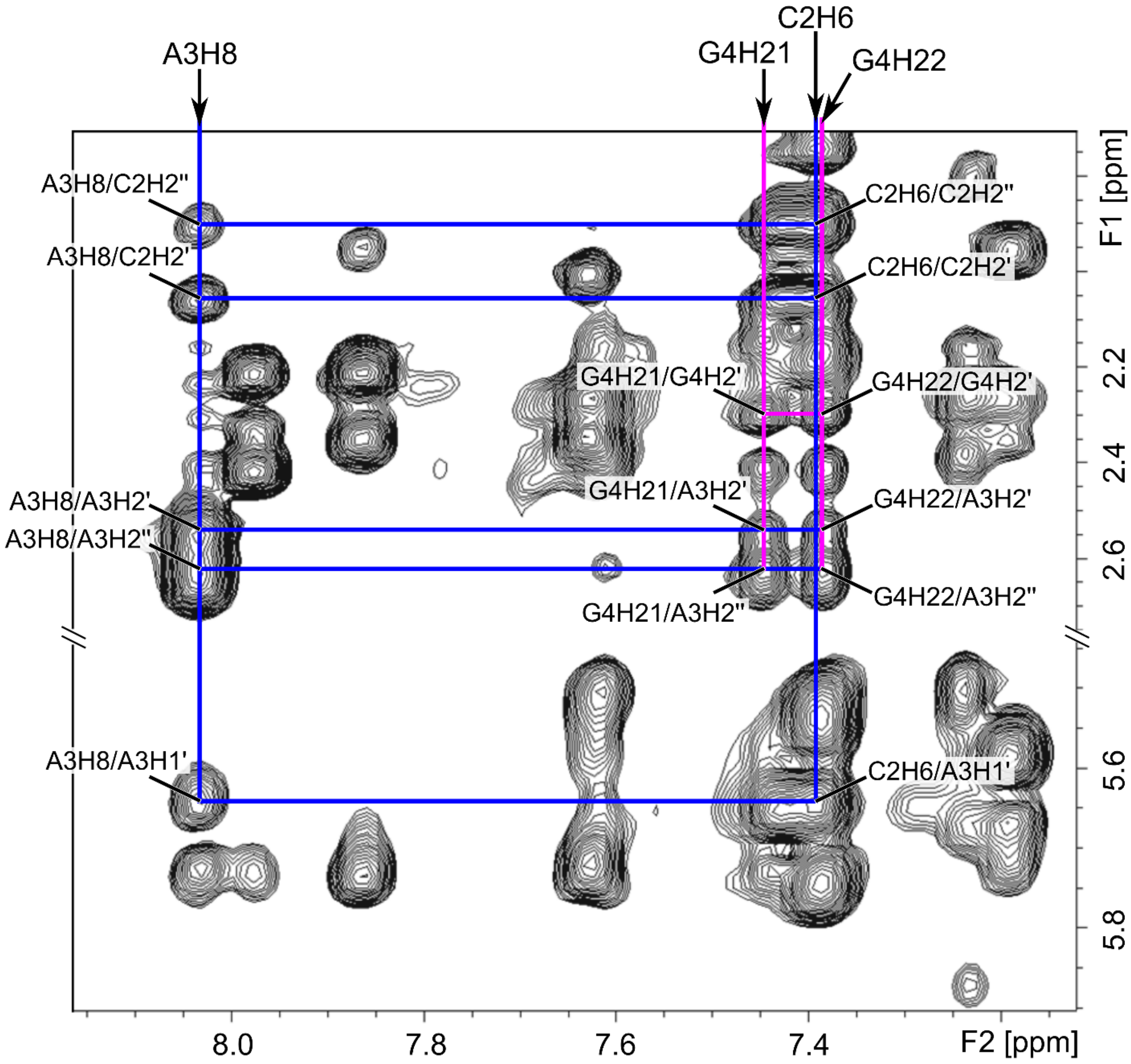

**Figure S5.** Regions of the 11-mer 2D-NOESY NMR spectra showing connectivity from C2 to G4. The C2H6/A3H1' and A3H8/A3H1' cross-peaks demonstrate the proximity between C2 and A3, whereas the G4H21/A3H2', G4H21/A3H2'', G4H22/A3H2', and G4H22/A3H2'' cross-peaks confirm the connectivity from A3 to G4.

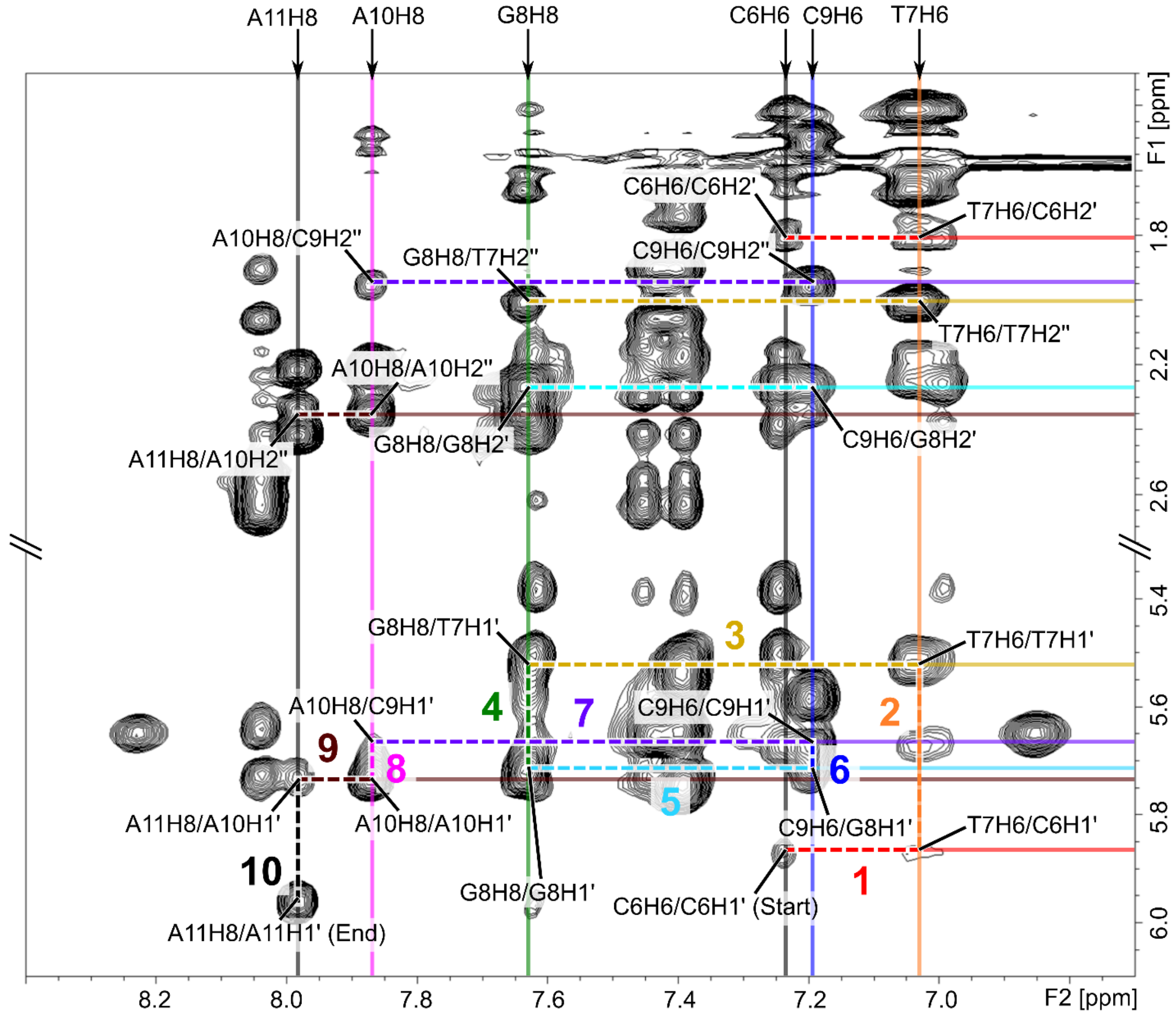

**Figure S6.** Sequential intra-strand connectivities for C6→T7→G8→C9→A10→A11 of the 11-mer. Cross-peaks in the non-exchangeable proton regions of the 2D-NOESY NMR spectra are labeled and indicated by intersecting lines. The sugar-to-base (H8/H6 to H1') connectivities are colored and numbered 1 – 10. NOEs to H2' or H2'' are indicated in the same color scheme and demonstrate internal consistency.

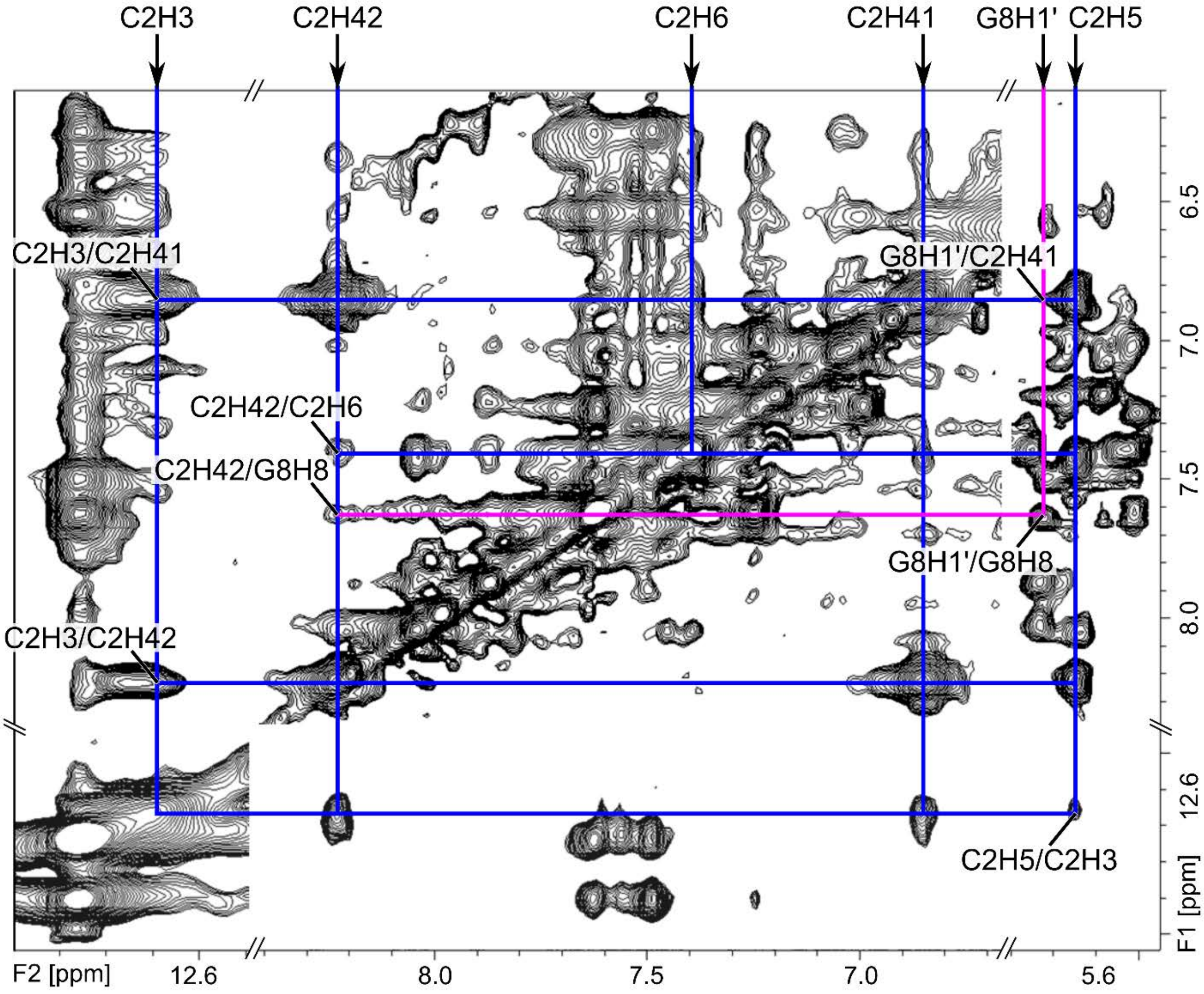

**Figure S7.** Regions of the 11-mer 2D-NOESY NMR spectra showing cross-peaks confirming the C2-C2$^+$ base pair. The C2H3/C2H41, C2H3/C2H42, and C2H5/C2H3 cross-peaks indicate internal consistency, whereas the C2H42/G8H8 and G8H1'/G8H8 cross-peaks confirm the proximity of C2H3 and G8.

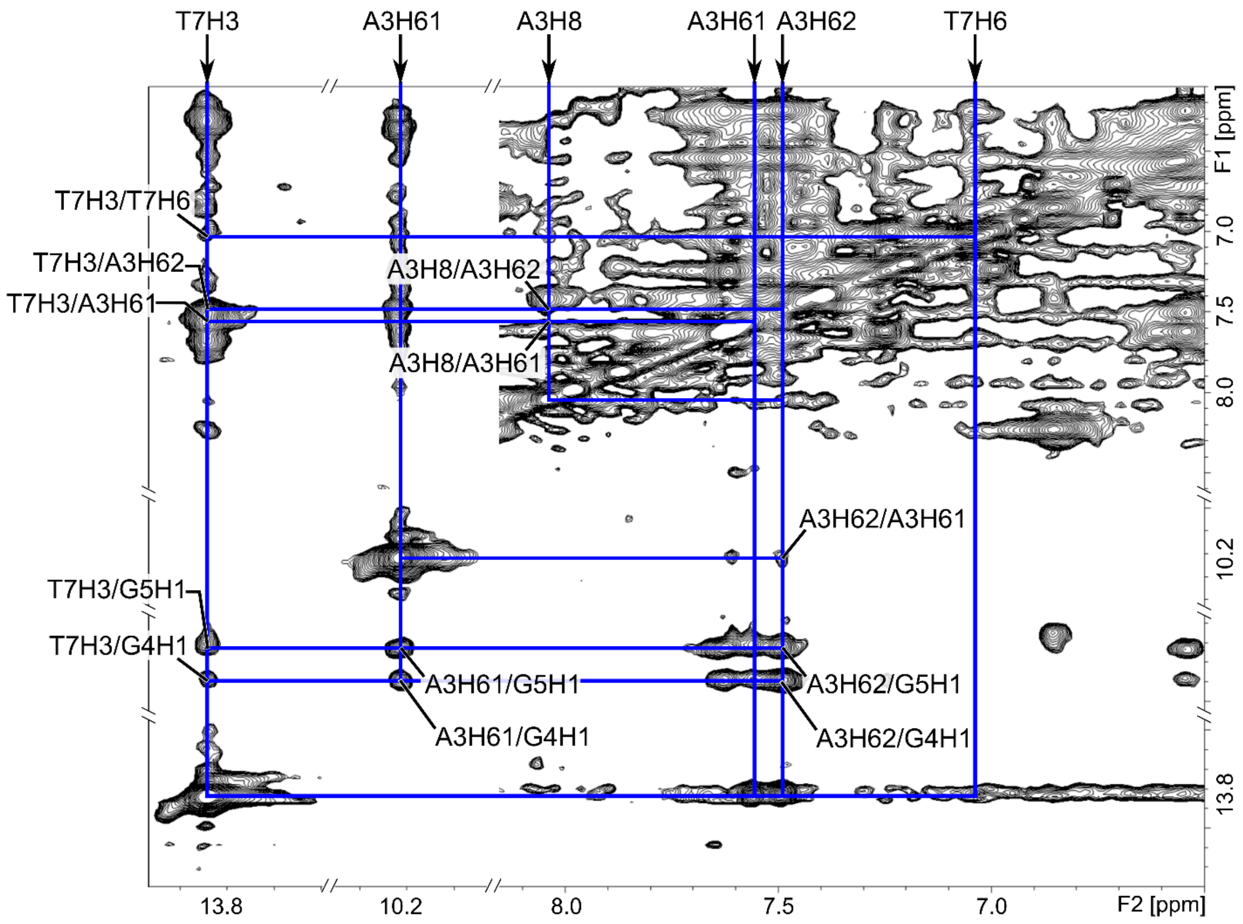

**Figure S8.** Regions of the 11-mer 2D-NOESY NMR spectra showing cross-peaks confirming the A-A-T base triple. The T7H3/A3H61 and T7H3/H62 cross-peaks confirm the T7-A3 base pair and the A3H8/A3H61, A3H8/A3H62, and A3H62/A3H61 cross-peaks from two independently assigned A3 residues provide evidence for the A3-A3 base pair. NOEs between the guanosine imino protons and the T7H3, A3H61, and A3H62 confirm the stacking between the base triple and G-tetrad.

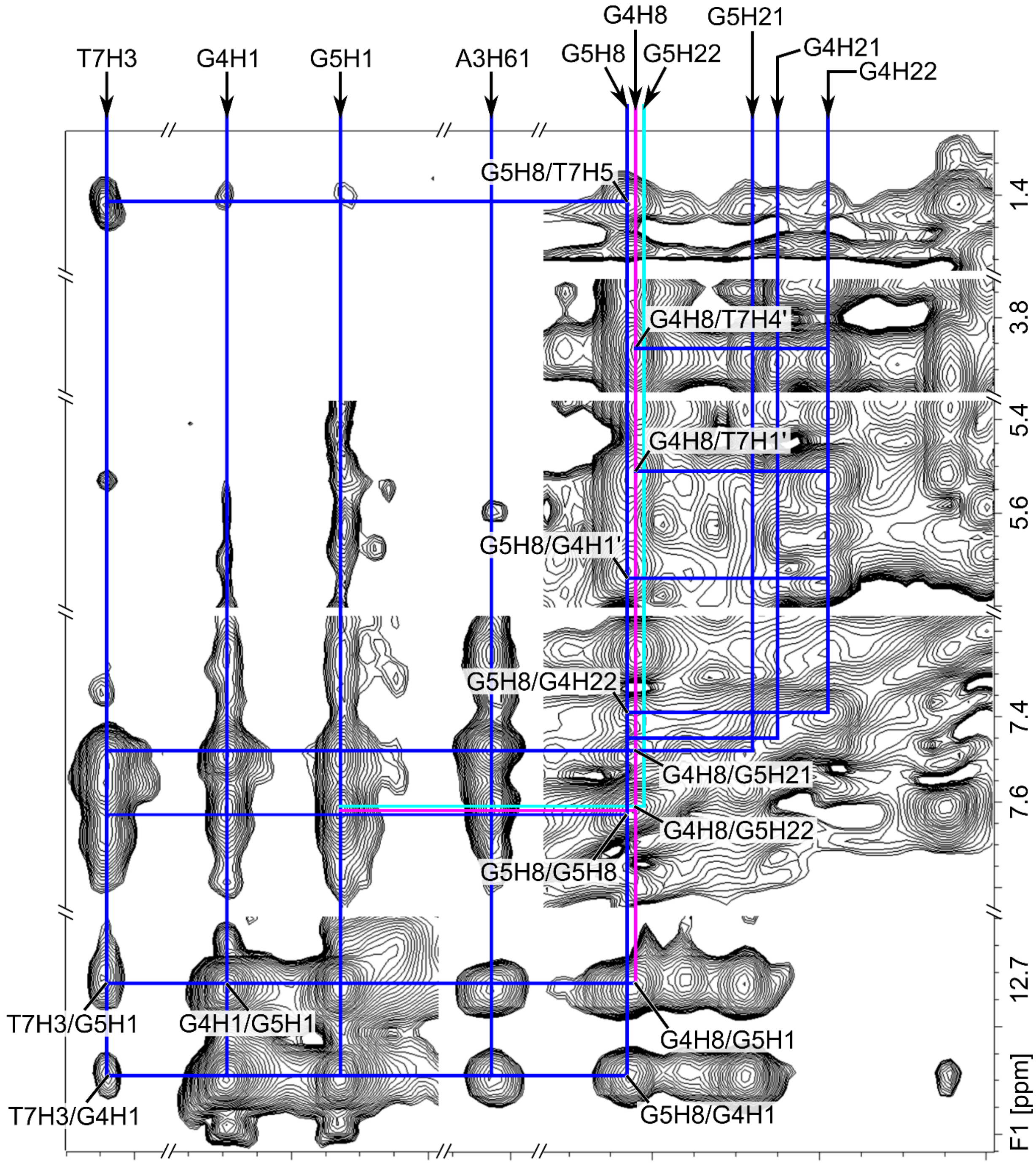

**Figure S9.** 2D-NOESY NMR spectra of the 11-mer showing cross-peaks confirming the G-tetrad. The G4H1/G5H1 cross-peak, in conjunction with the G4H8/G5H21, G4H8/G5H22, and G5H8/G4H22 cross-peaks, confirm hydrogen bonding between G4 and G5 through their Watson-Crick and Hoogsteen faces. The G5H8/T7H5 cross-peak, as well as resonances between G4H8 and the sugar protons of T7, confirm the arrangement of the guanosines in the G-tetrad in relation to the A-A-T base triple.

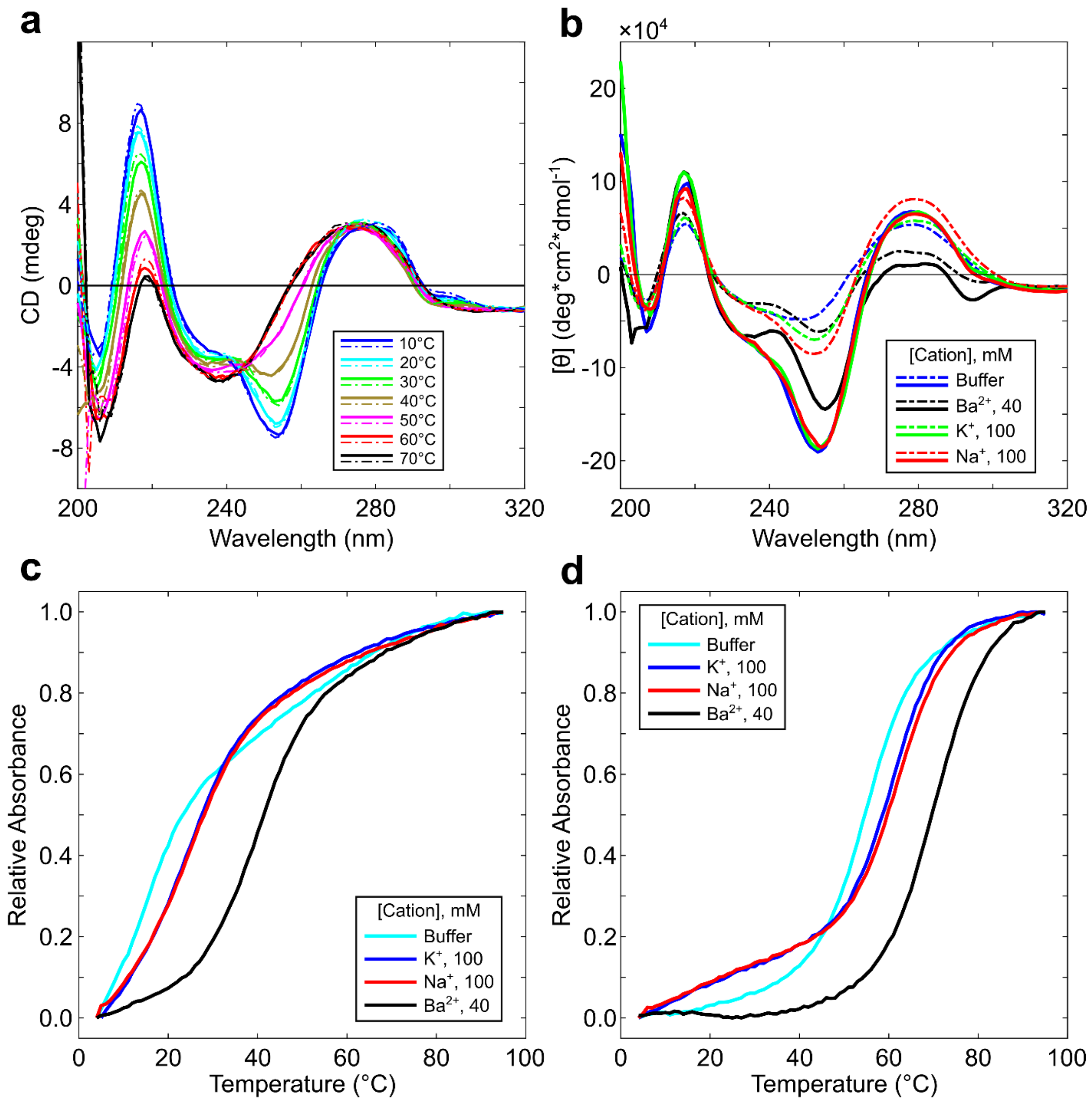

**Figure S10.** CD and Absorption spectra. (a) Forward (solid) and reverse (dash) CD melting curves of the 11-mer at 40 mM $Ba^{2+}$ collected at different temperatures. (b) CD spectra of the 11-mer (dotted) and 22-mer (solid) in sodium cacodylate buffer pH 6.0 or supplemented with 40 mM $Ba^{2+}$, 100 mM $K^+$, or 100 $Na^+$. Thermal denaturation curves from 4°C to 95°C of the 11-mer (c) or 22-mer.

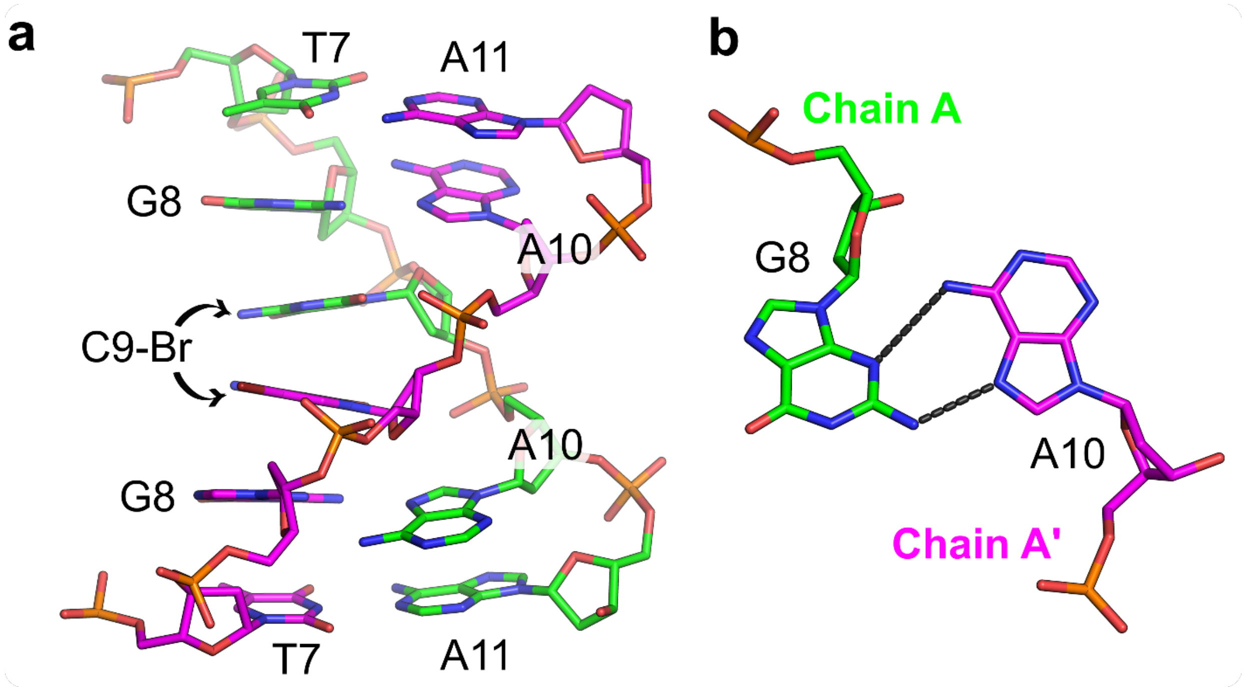

**Figure S11.** d(CCAGGCTGC$^{Br}$AA) Duplex. (a) Stick representation of the 3' antiparallel duplex formed from residues T7 through A11. (b) Residue G8 from Chain A (green) base pairs with A10 from Chain A' (magenta) through the G(N2)-A(N7) and G(N3)-A(N6) hydrogen bonds.

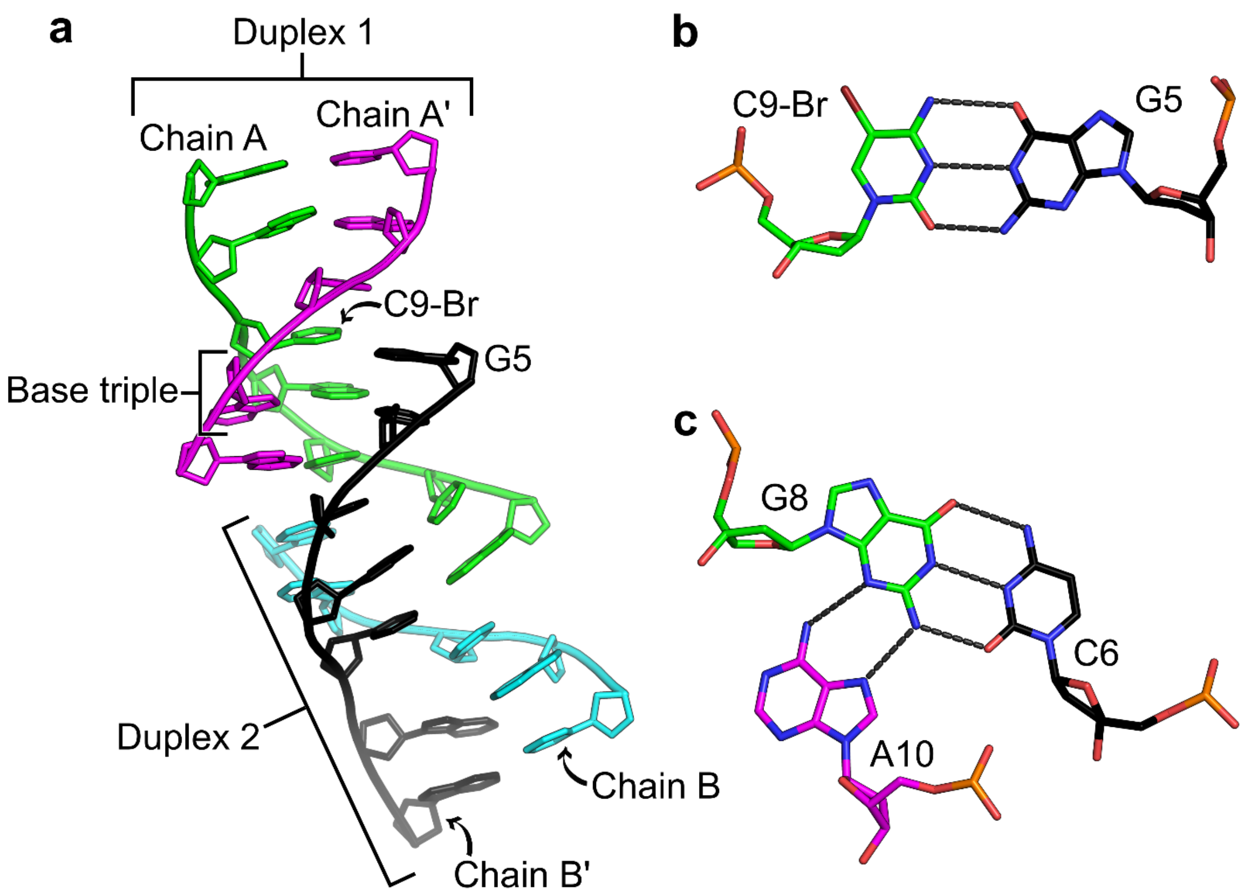

**Figure S12.** d(CCAGGC$^{Br}$TGCAA) Duplex Interactions. (a) Cartoon representation of the interactions between Duplex 1 (formed between Chains A and A') and Duplex 2 (formed between Chains B and B'). (b) Stick representation of the C9-Br-G5 base pair through standard Watson-Crick hydrogen bonding. (c) Stick representation of the C6-G8-A10 base triple. The G8-A10 base pair formed through G(N3)-A(N6) and G(N2)-A(N7) hydrogen bonding from Duplex 1 is converted into a base triple through Watson-Crick interactions with C6 from Duplex 2.
